# Deletion of autism risk gene Shank3 disrupts prefrontal connectivity

**DOI:** 10.1101/409284

**Authors:** Marco Pagani, Alice Bertero, Adam Liska, Alberto Galbusera, Mara Sabbioni, Maria Luisa Scattoni, Massimo Pasqualetti, Alessandro Gozzi

## Abstract

Mutations in the synaptic scaffolding protein Shank3 are a major cause of autism, and are associated with prominent intellectual and language deficits. However, the neural mechanisms whereby SHANK3 deficiency affects higher order socio-communicative functions remain unclear. Using high-resolution functional and structural MRI in mice, here we show that loss of *Shank3* (*Shank3B*^-/-^) results in disrupted local and long-range prefrontal functional connectivity, as well as fronto-striatal decoupling. We document that prefrontal hypo-connectivity is associated with reduced short-range cortical projections density, and reduced gray matter volume. Finally, we show that prefrontal disconnectivity is predictive of social communication deficits, as assessed with ultrasound vocalization recordings. Collectively, our results reveal a critical role of SHANK3 in the development of prefrontal anatomy and function, and suggest that SHANK3 deficiency may predispose to intellectual disability and socio-communicative impairments via dysregulation of higher-order cortical connectivity.

## Introduction

Mutations in genes coding for synaptic proteins are among the most well-characterized genetic deficits observed in individuals with autism spectrum disorders (ASD, de la Torre-Ubieta et al., 2016). The postsynaptic scaffolding protein SH3 and multiple ankyrin repeat domains 3 (SHANK3) is a critical orchestrator of macromolecular signaling complex at glutamatergic synapses (Peça et al., 2011; Tabet et al., 2017). In humans, full deletion of SHANK3 is found in a large fraction of Phelan-McDermid syndrome cases, a neurodevelopmental disorder characterized by ASD-like behaviors, intellectual disability (Phelan and McDermid, 2011) and profoundly impaired development of speech and language (Leblond et al., 2014; Monteiro and Feng, 2017). Recent genetic studies have also identified a large number of *Shank3* mutations in ASD patients not diagnosed with Phelan-McDermid syndrome, strongly implicating disruption or mutations of SHANK3 as one of the most frequent and penetrant monogenic cause of ASD and socio-communicative impairments (Leblond et al., 2014).

Animal studies have provided insight into the circuital dysfunctions produced by *Shank3* mutations. Prompted by the initial observation of self-injurious grooming in mice lacking the *Shank3B* isoform (Shank3B-/-, Peca et al., 2011), multiple investigations have probed basal ganglia and cortico-striatal function in mice harboring various *Shank3* mutations (Mei et al., 2016; Monteiro and Feng, 2017). These efforts have highlighted impaired synaptic structure and transmission and in striato-pallidal medium spiny neurons of SHANK3-deficient mice (Peça et al., 2011), an effect that is partially rescued by functional stimulation or adult re-expression of the protein in the affected neural populations (Mei et al., 2016; Wang et al., 2017). Functional deficits in cortico-striatal synchronization and excitability have also been described in Shank3 mutant mice (Peixoto et al., 2016; Wang et al., 2016). However, while these abnormalities serve as a plausible substrate for the motor stereotipies associated with *Shank3* disruption, the neural mechanism whereby SHANK3 deficiency affects higher order socio-communicative and cognitive functions remains unclear.

Systemic alterations in functional connectivity are prevalent in ASD (Di Martino et al., 2008) and have been regarded to account for the complex repertoire of symptoms exhibited by ASD patients (Vasa et al., 2016). In keeping with this notion, alterations in large-scale brain connectivity have been described in syndromic forms of autism, including *Cntnap2* polymorphism (Scott-Van Zeeland et al., 2010) and chromosome 16p11.2 microdeletion (Bertero et al., 2018). Based on these observations, here we hypothesize that SHANK3 insufficiency may impact higher-order socio-communicative functions via a dysregulation of inter-regional functional connectivity. To probe this hypothesis, we employed resting-state functional MRI (rsfMRI) (Gozzi and Schwarz, 2016), and retrograde viral tracing (Bertero et al., 2018) to map functional and structural connectivity in *Shank3B*^-/-^ mice. This mouse line lacks the exons encoding the PDZ domain of Shank3 (Monteiro and Feng, 2017), and exhibits ASD-related behavioral abnormalities, including robust repetitive behavior and social interaction deficits (Peça et al., 2011). We show that homozygous loss of *Shank3B* results in disrupted fronto-cortical and fronto-striatal connectivity, an effect associated with frontal gray matter hypotrophy and defective short-range cortico-cortical wiring. We also document that prefrontal connectivity deficits are predictive of socio-communicative impairments, as assessed with ultrasound vocalizations in a social interaction test. We suggest that *Shank3B* deletion may predispose to intellectual disability and socio-communicative impairments via a focal dysregulation of prefrontal connectivity.

## Results

### Disrupted prefrontal connectivity in *Shank3B null mice*

To identify regions exhibiting altered rsfMRI connectivity in *Shank3B*^-/-^ mice, we computed inter-group differences in long-range and local functional connectivity using spatially-unbiased aggregative network-based metrics (Liska et al., 2017). These analyses revealed a robust reduction in long-range and local connectivity in prefrontal cortical areas of *Shank3B*^*-/-*^ mutants (*t*-test, *p* < 0.01 FWE cluster-corrected, with cluster defining threshold of |*t*| > 2, [*p* < 0.05], Fig. 1A-C). Long-range connectivity reductions appeared to be more widespread than corresponding local connectivity deficits, and included retrosplenial and prefrontal-infralimbic cortices, as well as bilateral striatal areas and anterior insular cortices (Fig. 1A). *Posthoc* region-based quantifications confirmed these effects (Fig. 1B-D, *t*-test, *p* < 0.05). To corroborate these findings and probe long-range targets of the observed functional hypoconnectivity, we next calculated genotype differences in inter-regional rsfMRI connectivity by applying network-based statistics (NBS) to a whole-brain network parcellation (Zalesky et al., 2010). This analysis confirmed the presence of predominant regional hypo-connectivity in prefrontal regions and cingulate-retrosplenial midline areas (*t*-test, link-wise threshold *t* > 3.5 [*p* < 0.001], FWE corrected at *p* < 0.05, Fig. 1E-F). It also revealed the presence of long range desynchronization between prefrontal regions and other long-range cortical targets, including entorhinal, piriform and motor cortices (Fig. 1E-F). These results reveal the presence of impaired functional synchronization in prefrontal cortical areas of *Shank3B* ^-/-^mice.

**Figure 1.**
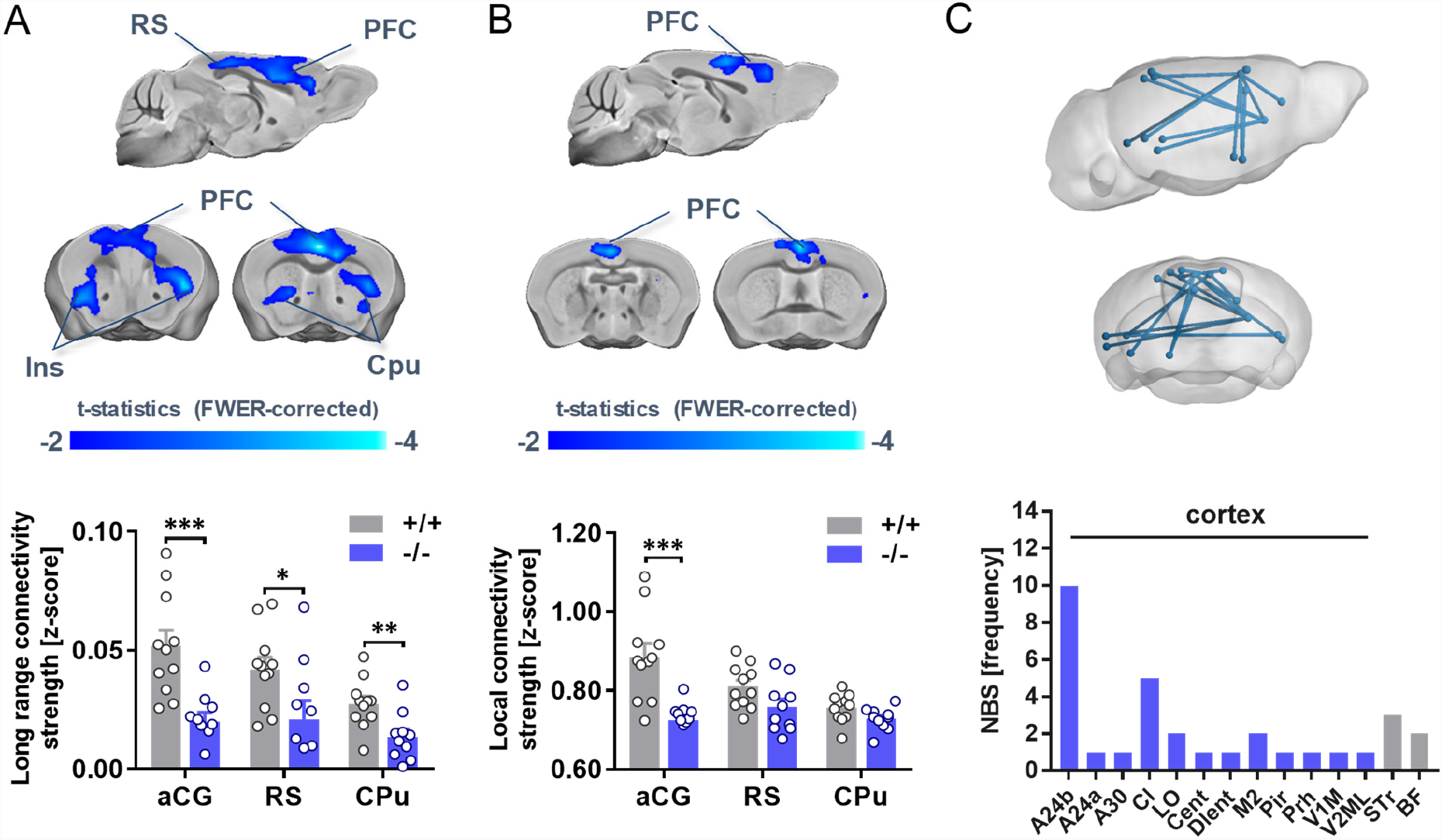
Reduced local and long-range connectivity in prefrontal areas of *Shank3B* ^-/-^ mice. Reduced long-range (A) and local (B) connectivity in the prefrontal cortex of *Shank3B*^-/-^ mutants (blue, *t* > 2, *p* < 0.01 FWE cluster corrected). Foci of reduced long-range connectivity were also present in the retrosplenial and insular cortex, and striatal areas. (*p<0.05, **p<0.01, ***p<0.01, Student’s *t*-test). (C) Inter-group comparison of rsfMRI connectivity in regionally-parcellated brains using network based statistics (NBS). The plot shows, for each region, the number of links exhibiting reduced rsfMRI connectivity identified with NBS (*t* > 3.5, FWE corrected at *p* < 0.05). CPu: caudate putamen; PFC: prefrontal cortex; RS: retrosplenial cortex.

### Reduced default-mode-network and fronto-striatal connectivity in *Shank3B*^-/-^ mice

Long-range connectivity mapping revealed foci of weaker synchronization in prefrontal and retrosplenial cortex of *Shank3B*^-/-^ mice, an ensemble of associative regions that act as pivotal nodes of the mouse default mode network (DMN) (Sforazzini et al., 2014). To probe the functional integrity of this network, we performed a seed-based correlation analysis along its midline extension. In keeping with global connectivity mapping, we observed foci of reduced connectivity throughout the antero-posterior cingulate-retrosplenial axis of the DMN (*t*-test, *p* < 0.01 FWE cluster-corrected, with cluster defining threshold of |*t*| > 2 [*p* < 0.05]), as well as reduced involvement of temporal cortical areas in *Shank3B*^-/-^ mice (Fig. 2A). A quantification of DMN connectivity via multiple prefrontal-DMN seed pairs (Fig. 2B) revealed a clear disconnection of the prefrontal cortex with the rest of this network (two-way repeated-measures ANOVA, genotype effect, *F*_1,19_ = 15.01, *p* = 0.001, Fig. 2B). Similarly, seed-based mapping of the anterior caudate-putamen, a region exhibiting focal under-connectivity (Fig. 1A), revealed genotype-dependent functional desynchronization between basal ganglia and the anterior cingulate cortex (Fig. 2C-D, *t*-test, genotype effect, *t*_19_ = 2.30, *p* = 0.03, Fig. 2D). Striatal regions also exhibited robustly decreased inter-hemispheric connectivity (*t*-test, genotype effect, *t*_19_ = 2.90 *p* = 0.009, Fig. S1). Inter-hemispheric connectivity between motor-sensory cortical networks was otherwise largely unimpaired, supporting the network specificity of these findings (Fig. S1). Interestingly, we found *Shank3B* ^-/-^ mutants to exhibit decreased inter-subject similarity as assessed with group-based homotopic patterns (Fig. S2A-B), a finding recapitulating idiosyncratic inter-hemispheric connectivity patterns in human ASD patients (Hahamy et al., 2015). This effect was especially prominent in cortical regions (Fig. S2C-D).

**Figure 2.**
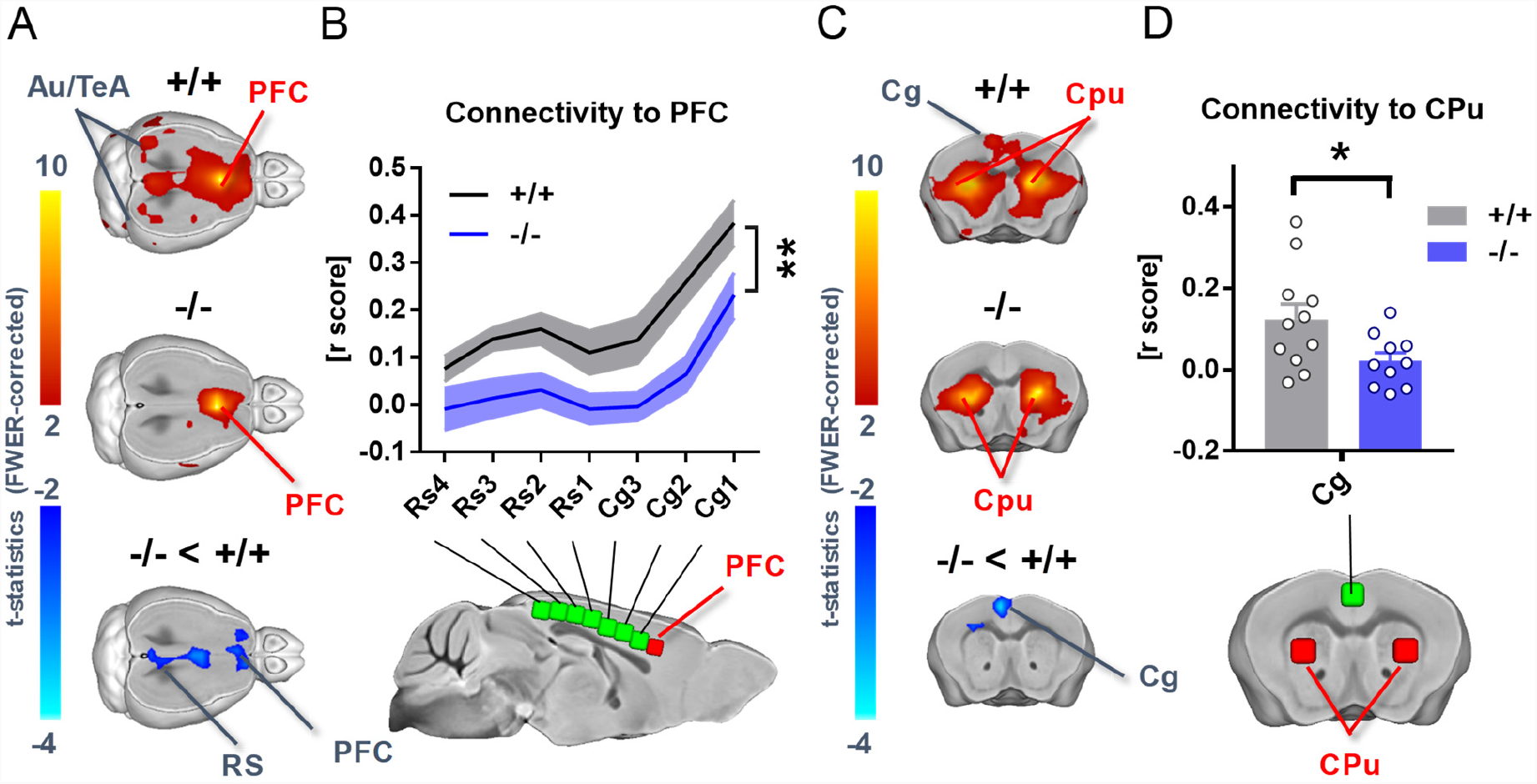
DMN and fronto-striatal under-connectivity in *Shank3B*^-/-^ mice. (A) Reduced rsfMRI connectivity in the midline component of DMN between the anterior cingulate and the retrosplenial cortex of *Shank3B*^-/-^ mice compared to wild-type littermates (blue/light blue, *t* > 2, *p* < 0.01 FWE cluster corrected). (B) PFC-to-seed rsfMRI connectivity profiling corroborates reduced functional coupling between prefrontal cortex and cingulate-retrosplenial component of DMN (2-way repeated-measures ANOVA, genotype effect, *F*_1,19_ = 15.01, *p* = 0.001). (C) Seed-based analysis of the striatum highlighted a reduction in fronto-striatal connectivity. (D) Fronto-striatal de-synchronization as assessed using a reference volume of interest (VOI) in the anterior cingulate cortex (t_19_ = 2.30, *p* = 0.03, D). Red/yellow indicates positive correlation with seed regions (*t* > 2, *p* < 0.01 FWE cluster corrected). Blue indicates foci of significantly reduced connectivity in *Shank3B*^-/-^ mutants (*t* > 2, *p* < 0.01 FWE cluster corrected). VOIs location are depicted in red, seeds in green. Au: auditory cortex; Cg: cingulate cortex; CPu: striatum; PFC: prefrontal cortex; Rs: retrosplenial cortex; TeA: temporal associative cortex. *p<0.05, **p<0.01.

Importantly, no genotype-dependent differences in anesthesia sensitivity were detected when assessed with mean arterial blood pressure mapping (*t*-test, *t*_19_=1.23 *p* = 0.23; Fig. S3A), amplitude of cortical BOLD signal fluctuations (*t*-test, *t*_19_ = 1.15, *p* = 0.26; Fig. S3B) and minimal alveolar concentration of anesthetic (*t*-test, *t*_19_ = 1.02, *p* = 0.31; Fig. S3C), three physiological measures sensitive to anesthesia depth (Steffey et al., 2003; Liu et al., 2011; Bertero et al., 2018). Together with the observation of region-dependent rsfMRI alterations, these findings argue against a significant confounding contribution of anesthesia to the observed hypo-connectivity phenotype.

### Prefrontal under-connectivity in *Shank3B*^*-/-*^ predicts socio-communicative deficits

Shank3 deficiency in humans is associated with marked language and communicative deficits. To test whether Shank3 mutations can similarly affect socio-communicative function in rodents and identify the neural circuits responsible for this phenomenon, we measured sociability and ultrasound vocalizations in a male-female interaction test and we correlated each of these measurements with rsfMRI connectivity metrics we previously recorded in the same subjects. Behavioral testing revealed unaltered total sniffing and social communication in *Shank3B*^-/-^ mice, along with a trend for reduced sociability as assessed with anogenital sniffing duration (*t*-test, *t*_19_ = 2.21, *p* = 0.04, Fig. S4). The same recordings revealed a marked reduction in the frequency of ultrasound vocalizations in *Shank3B*^*-/-*^ mice (*t*-test, *t*_19_ = 3.60, *p* = 0.002, Fig. 3A). Interestingly, voxel-wise correlation mapping revealed that DMN hypo-connectivity was significantly associated with reduced vocalizations, with evidence of robust correlation in prefrontal and cingulate cortices (*t*-test, *p* < 0.01 FWE cluster-corrected, with cluster defining threshold of |*t*| > 2 [*p* < 0.05], Fig. 3B). These findings suggest that prefrontal under-connectivity is predictive of socio-communicative impairments observed in these mice. A regional quantification of this effect in the prefrontal cortex confirmed a strong positive correlation between these two readouts (*r* = 0.67, *p* < 0.001, Fig. 3C). Voxel-wise correlation mapping did not reveal any significant association between ultrasound vocalization frequency and cortico-striatal rsfMRI connectivity (*t*-test, *p* > 0.01 FWE cluster-corrected, with cluster defining threshold of |t| > 2 [p < 0.05]), corroborating the circuital specificity of our findings. These results point at prefrontal connectivity deficits as a putative network substrate for the socio-communicative deficits exhibited by *Shank3B*^*-/-*^ mice.

**Figure 3.**
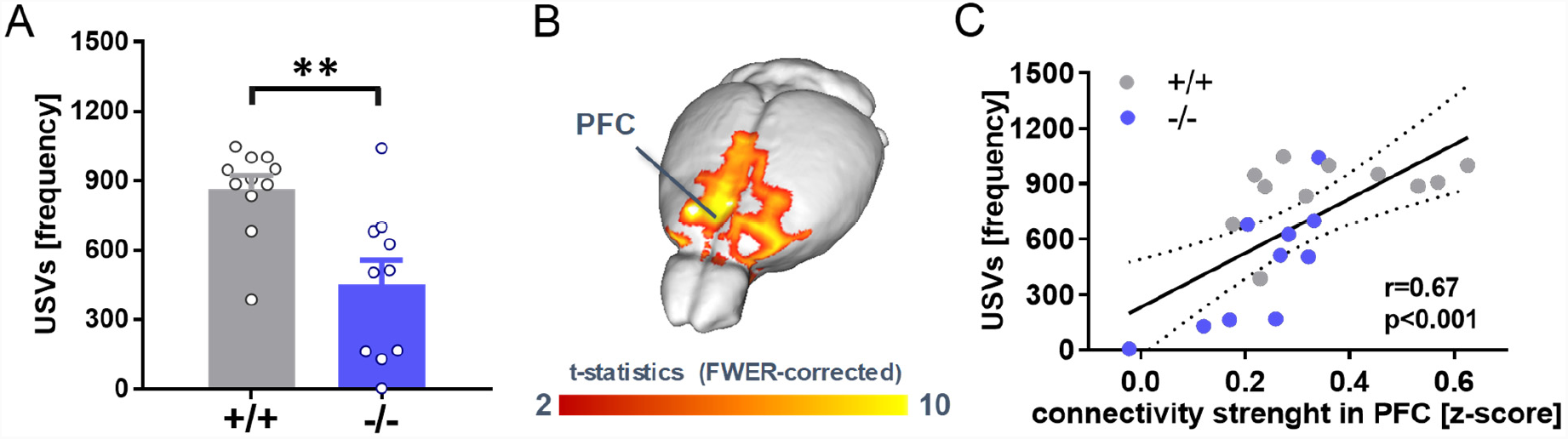
Prefrontal connectivity predicts socio-communicative dysfunction. (A) Reduced ultrasound vocalization frequency in *Shank3B*^-/-^ mice (*t*-test, *t*_24_ = 2.88, *p* = 0.008). (B) Voxel-wise correlation mapping revealed a significant positive correlation between prefrontal rsfMRI connectivity and vocalization frequency (*t*-test, *p* < 0.01 FWE cluster-corrected, with cluster defining threshold of |*t*| > 2 [*p* < 0.05]). (C) Quantification of this relationship in the prefrontal cortex (*r* = 0.67, *p* < 0.001, *n* = 21). \*\**p* < 0.01.

### Reduced prefrontal gray matter volume in *Shank3B*-/-mice

To investigate whether *Shank3B* deficiency in mice produces brain-wide cortical neuro-anatomical alterations, we used high-resolution structural MRI to obtain spatially-unbiased maps of gray matter volume with voxel-resolution (Pagani et al., 2016a). Notably, inter-group comparisons revealed extended reductions in cortical gray matter in *Shank3B*^-/-^ mice, encompassing a network of cortical regions exhibiting remarkable neuroanatomical overlap with prefrontal, midline and temporal nodes of the mouse DMN (*t*-test, *p* < 0.01 FWE cluster-corrected defining threshold of |*t*| > 2 [*p* < 0.05], Fig. 4A). Atlas-based volumetric estimation of regional gray matter volume confirmed the presence of reduced gray matter volume in cortical constituents of the mouse DMN (*t*-test, *p* < 0.05, Fig. 4B). Interestingly, we found prefrontal gray matter volume to be significantly correlated with long-range, but not-short range, connectivity strength (Fig. 4C, r = 0.592, *p* = 0.009, and r = 0.119, *p* = 0.636, respectively), establishing a putative link between these two findings.

**Figure 4.**
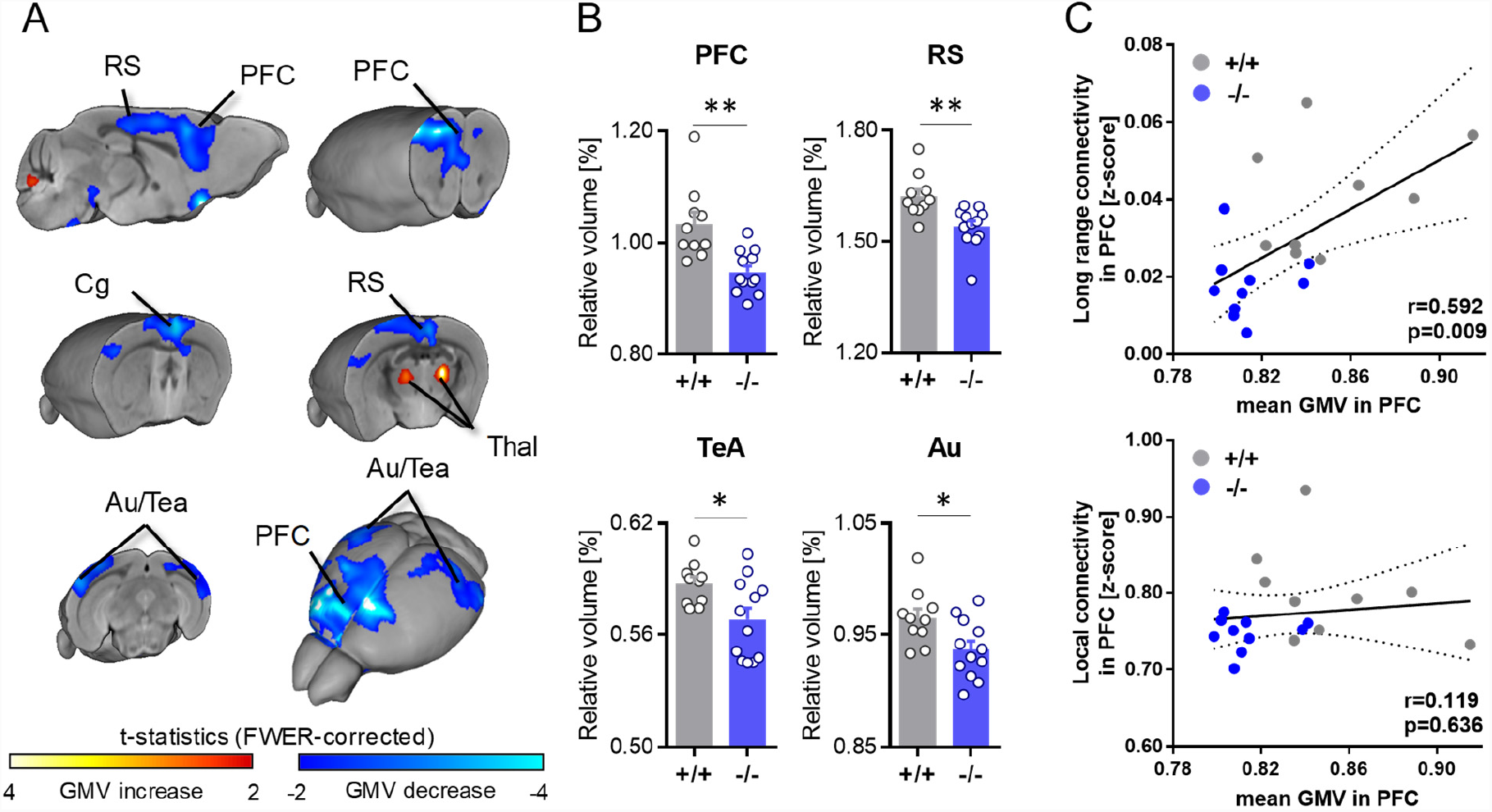
Reduced gray matter volume in prefrontal regions of *Shank3B* ^-/-^ mice. (A) Structural MRI revealed a prominent reduction of gray matter volume (GM) volume in prefrontal and retrosplenial cortices of *Shank3B*^-/-^ mutants. Statistically significant volume reductions were also apparent in peri-hippocampal associative areas of *Shank3B*^-/-^ mice (*t*-test, *p* < 0.01 FWE cluster-corrected, with cluster defining threshold of |*t*| > 2 [*p* < 0.05]). (B) Regional volumetric analysis confirmed the results of voxel-wise statistical mapping. (C) Pearson’s correlation between long range and local rsfMRI connectivity, and gray matter volumes. A positive correlation between prefrontal long-range connectivity and gray matter was observed in prefrontal areas (*r* = 0.592, *p* = 0.009, n = 19). Au: auditory cortex; Cg: cingulate cortex; PFC: prefrontal cortex; RS: retrosplenial cortex; TeA: temporal association area; Thal: thalamus. \**p* < 0.05, \*\**p* < 0.01, Student’s *t*-tests.

### Reduced short-range projection density in *Shank3B*-/-mice

Recent studies have revealed mesoscale wiring alterations as a possible correlate for long-range functional desynchronization in genetic models of autism (Liska et al., 2017; Bertero et al., 2018). To investigate the presence of neuroanatomical alterations in prefrontal areas of *Shank3B* mutant mice, we first examined whether these alterations could be related to differences in neuronal density, similar to findings recently described in a primate model of Shank-deficiency (Zhao et al., 2017). Post-mortem quantification of neuronal somas in the medial prefrontal cortex did not reveal genotype-dependent differences in NeuN-positive cell density (*Shank3B*^*+/+*^: 0.439± 0.061 cells/100pixel; *Shank3B*^*-/-*^: 0.485 ± 0.062 cells/100pixel; *p* = 0.27, two tailed *t*-test).

We next carried out monosynaptic retrograde tracing of the prefrontal cortex using a recombinant retrograde rabies virus. We then quantified the fraction of retrogradely labeled cells within prefrontal areas (Fig. 5A) exhibiting significant local rsfMRI hypo-connectivity (Fig. 5B), and in a representative set of distal source regions (Fig. 5C). Quantifications of labeled cells within the prefrontal foci of hypoconnectivity, revealed significantly reduced number of locally projecting neurons in *Shank3B* mutants (*t*-test, *t*_7_ = 3.90, *p* = 0.025, Fig. 5B-C). No inter-group difference in the number of prefrontal-projecting neurons was observed in any of the long-range source regions (Fig. 5C, *p* > 0.05, all regions, FDR corrected). These results reveal a possible contribution of defective short-range mesoscale axonal wiring to the observed prefrontal under-connectivity.

**Figure 5.**
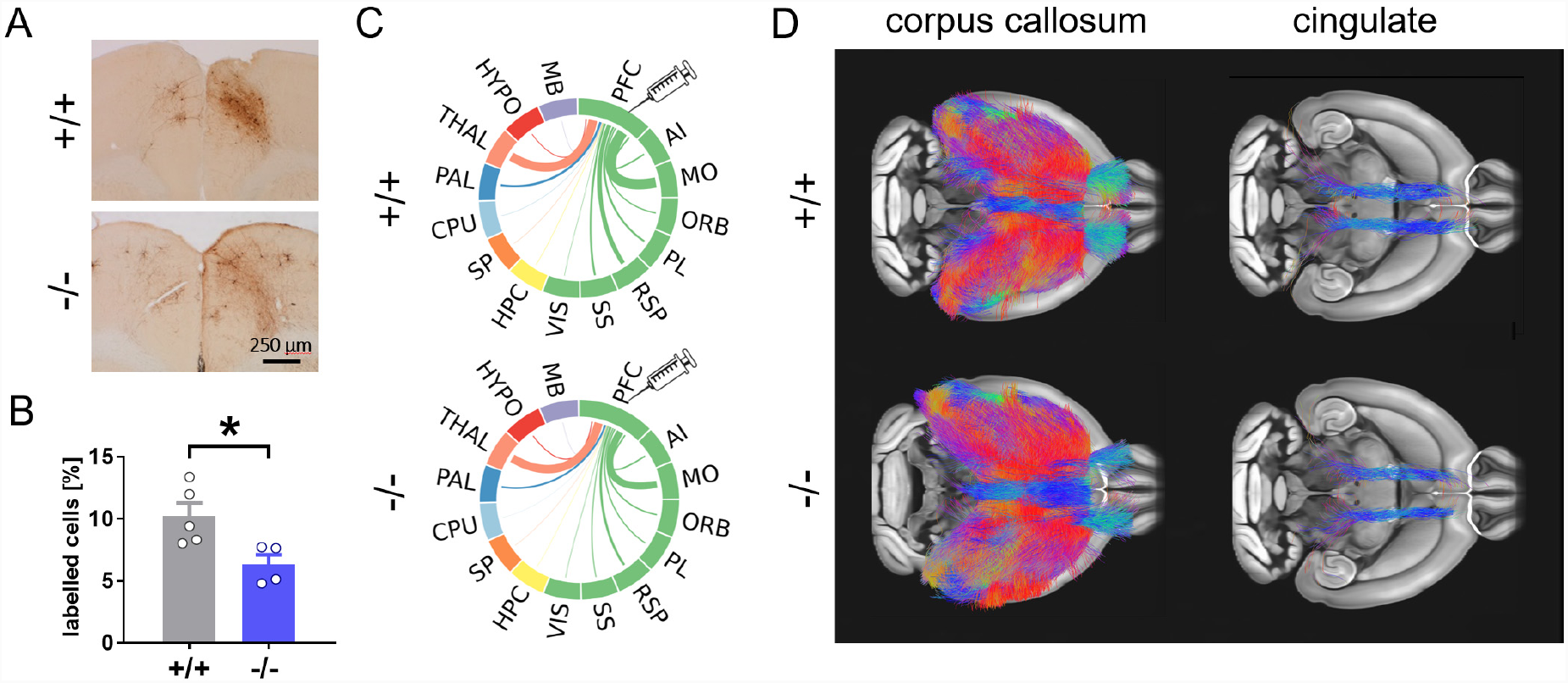
Neural rewiring in the prefrontal cortex of *Shank3B*^-/-^ mice. (A) Enlarged view of the distribution of retrogradely labelled cells in representative coronal sections of the cingulate cortex (bregma 1.25mm) exhibiting decreased number of retrogradely labelled cells in Shank3B**^-/-^** mice. Scale bar: 250 μm. (B) Regional quantification of the relative regional number of retrogradely labelled cells in prefrontal areas exhibiting reduced *local* rsfMRI functional connectivity (\**p* < 0.05, Student’s *t*-test, means ± SEM). (C) Circular layout showing the relative number of retrogradely labelled cells. The thickness of links is proportional to the relative number of labelled cells. Cortical areas are shown in green. (D) No apparent genotype-dependent alterations in white matter fiber-organization were observed with tractography when we dissected corpus callosum (left) and cingulate (right). PFC: prefrontal cortex; ACA: anterior cingulate cortex; AI: agranular insular cortex; MO: motor cortex; ORB: orbital cortex; PL: prelimbic cortex; RSP: retrosplenial cortex; SS: somatosensory cortex; VIS: visual cortex; HPC: hippocampus; SP: cortical subplate; CPU: striatum; PAL: globus pallidus; Thal: thalamus; HYPO: hypothalamus; MB: midbrain.

We next probed overall white matter microstructural integrity using diffusion-weighted MRI, and carried out *ex vivo* tractography to virtually dissect the corpus callosum and cingulum, two major fiber bundles linking interhemispheric and antero-posterior cortical areas. No genotype-dependent white matter reorganization (*p* > 0.05, Fig. 5D) or microstructural alterations (fractional anisotropy, median diffusivity, radial diffusivity, *p* > 0.05, all parameters voxel-wise) were observed. These findings argue against a contribution of gross white matter reorganization to the observed functional disconnectivity.

## Discussion

Here we document disrupted fronto-cortical functional connectivity in mice lacking autism-risk gene *Shank3,* and show that this trait is predictive of impaired socio-communicative functions. We also describe reduced fronto-cortical volume and short-range neuronal wiring as plausible mesoscopic correlates of the observed disconnectivity. These findings establish a link between a prevalent monogenic form of ASD and aberrant neocortical connectivity, suggesting that Shank3 deficiency can predispose to socio-communicative dysfunction and intellectual disability via dysregulation of higher-order cortical synchronization.

Disruption in large-scale brain network activity has been consistently observed in individuals with autism, and is regarded to partially account for language and cognitive impairments manifested in ASD (Vasa et al., 2016). However, the clinical heterogeneity of ASD has so far precluded a systematic and reliable identification of network alterations specific to distinctive pathophysiological or genetic etiologies. Our results suggest that Shank3 deficiency, one of the most common genetic alterations in ASD patients, leads to selective desynchronization of associative and integrative cortical regions. This finding adds to recent human and animal evidence implicating disrupted functional connectivity as a network-level alteration of pathophysiological relevance for ASD, a feature that has been observed in an increasing number of syndromic forms of ASD, such as cortical dysplasia produced by CNTNAP2 (Liska et al., 2017), 16p11.2 microdeletion (Bertero et al., 2018), haploinsufficiency in chromatin domain 8 protein (Suetterlin et al., 2018) and genetic variants in Met receptor tyrosine kinase (Rudie et al., 2012).

The observation of reduced prefrontal connectivity in *Shank3B*^-/-^ mice opens the way to targeted investigations of the specific neural brain mechanisms underlying this functional impairment, and its significance for the manifestation of ASD symptoms. Given the critical contribution of fast-spiking interneurons in orchestrating long-range functional synchronization (Sohal et al., 2009), the presence of parvalbumin–positive interneuron dysfunction in *Shank3B*^*-/-*^ mice (Gogolla et al., 2014; Filice et al., 2016) serves as a putative neurophysiological correlate for the observed fMRI desynchronization. This hypothesis is consistent with our observation of similarly-impaired long-range prefrontal connectivity in *Cntanp2*-null mice, a mouse line exhibiting ASD-like behaviors and characterized by reduced GABA interneuron density (Liska et al., 2017). Interestingly, as observed in *Cntnap2* mutants, *Shank3B*^-/-^ mice also revealed mesoscale wiring defects involving a reduced projection density of cortico-cortical pyramidal neurons. While the exact mechanisms leading to improper cortico-frontal circuit assembly remain to be firmly established, it has been recently shown that interneuron-dependent homeostatic mechanisms taking place during development can critically sculpt cortical networks (Wong et al., 2018), hence putatively implicating dysfunctional GABAergic maturation in the defective wiring observed in these two genetic models of ASD. Such developmental mechanisms, together with imbalances in the excitatory output of specific regional districts (Peixoto et al., 2016), could conceivably lead to permanent imbalances in inter-regional connectivity and synchronization. The presence **o**f reduced gray matter volume in the same prefrontal regions exhibiting dysfunctional coupling recapitulates neuroimaging observation in ASD patient cohorts, in which reduced gray matter volume in the temporal lobe was associated with locally decreased functional connectivity (Mueller et al., 2013). This abnormality does not appear to reflect genotype-dependent differences in neuronal density or major histological alterations in the PFC of SHANK3-deficient mice, and are therefore more likely to reflect a reduced functional engagement of these cortical substrates as a result of their abnormal connectivity. In keeping with this notion, morphological investigations of gray matter volume changes detected by MRI in rodents have revealed a direct link between imaging findings and dendritic spine density, but not neuronal density (Keifer Jr et al., 2015). Based on these results, the observed reduction in gray matter volume could tentatively be linked to the reduced dendritic spine density previously reported in different mouse models of Shank3 deficiency (Peca et al., 2011; Wang et al., 2016). Future investigations of the microstructure of the PFC with respect to other regional districts may be informative as to the nature of this macroscopic gray matter abnormality.

The presence of functional prefrontal alterations in *Shank3*-deficient mutants extends our knowledge of the neural circuitry disrupted by mutations in this autism-risk gene, and shifts the focus from the basal ganglia, in which the role of Shank3 deficiency has been compellingly elucidated (Peca et al., 2011; Peixoto et al., 2016), to integrative higher-order cortical regions of relevance for ASD. The role of the prefrontal cortex as a crucial mediator of social recognition in rodents has prompted recent investigations of cortical dysfunction in Shank3-deficient mice, with initial results that are in excellent agreement with our findings. For example, Duffney et al., (2015) recently demonstrated that reduced social preference in Shank3^+/ΔC^ mice is associated with diminished NMDA receptor function and distribution in prefrontal regions. In the same mouse model, social deficits were ameliorated by treatment with a histone deacetylase (HDAC) inhibitor, highlighting a contribution of epigenetic mechanisms to the expression of social deficits in Shank3-deficiency (Qin et al., 2018). Our results are in keeping with these observations, and suggest that the prefrontal dysfunction that characterizes SHANK3-deficient mice affects complex socio-communicative behavior via a large-scale involvement of distributed fronto-striatal and fronto-cortical substrates. Notably, we found the latter connectional deficits to be predictive of socio-communicative function as assessed with ultrasound vocalizations during a social interaction task. While only correlative, this finding pinpoints a putative large-scale circuit for the expression of social communication, and strengthens translational research of Shank3 deficiency by showing that communication deficits, one of the foremost disabilities observed in Phelan Mc-Dermid syndrome and related shankopathies, can be recapitulated in rodent models. Interestingly, studies in rodents have shown a critical role of motor-sensory cortical areas in the expression of distress-related vocalizations, a set of regions that however appears to exhibit unaltered patterns of functional connectivity in our study (Sia et al., 2013). These results suggest that the communication deficits produced by SHANK3-deficiency are a top-down consequence of impaired social recognition, as opposed to primary dysfunction of motor-sensory areas related to the expression of vocalizations.

The use of directly translatable measurements of brain function and anatomy represents a distinctive feature of our approach that can be leveraged to cross-fertilize clinical and preclinical investigations of the macroscale substrates affected by ASD-related etiologies. Mouse models recapitulating syndromic forms of human disease-causing mutations can be used to establish causal relations between ASD-related genetic etiologies, abnormal macroscale connectivity and behavioral correlates, thus shedding light on the neurobiological mechanisms underlying ASD (Liska and Gozzi, 2016). Notably, using the same imaging paradigm employed here, we recently showed that orthologous 16p11.2 deletion results in remarkably similar patterns of functional under-connectivity in humans and mice, lending support to the possibility of directly relating neuroimaging readouts across species (Bertero et al., 2018). As neuroimaging data in genetically characterized patient populations are becoming increasingly available (Feliciano et al., 2018), our findings will serve as a translational benchmark to assess the mechanistic relevance and predictive validity of the use of rodents to model the network disruption produced by ASD-related etiologies.

In conclusion, our work shows that SHANK3-deficiency leads to disrupted prefrontal functional connectivity, an effect associated with aberrant neuronal wiring, reduced fronto-cortical gray matter volume, and impaired social communications. Our findings suggest that *Shank3* deletion may predispose to neurodevelopmental disorders and autism through dysregulation of connectivity in higher-order cortical areas, and provide a translational model for investigating connectional perturbations associated with ASD and related developmental disorders.

## Acknowledgments

The study was funded by grants from the Simons Foundation (SFARI 314688 and 400101, A. Gozzi). A. Gozzi also acknowledges funding from the Brain and Behavior Foundation (2017 NARSAD independent Investigator Grant).

## Material and methods

### Ethical statement

Animal studies were conducted in accordance with the Italian Law (DL 26/2014, EU 63/2010, Ministero della Sanità, Roma) and the recommendations in the Guide for the Care and Use of Laboratory Animals of the National Institutes of Health. All surgical procedures were performed under anesthesia.

### Resting-State fMRI

Male *Shank3B*^-/-^ (*n=* 10, 19-21 weeks old) and age-matched control *Shank3B*^+/+^ littermates (*n=*11) were housed under controlled temperature (21 ± 1 °C) and humidity (60 ± 10%). Resting-state functional MRI (rsfMRI) data were recorded as previously described (Ferrari et al., 2012; Sforazzini et al., 2016). Briefly, animals were anaesthetized with isoflurane (5% induction), intubated and artificially ventilated (2% maintenance). The left femoral artery was cannulated for continuous blood pressure monitoring throughout imaging sessions and for arterial blood sampling. After surgery, isoflurane was discontinued and replaced with halothane (0.7%). Functional data acquisition commenced 45 min after isoflurane cessation. Arterial blood gases (paCO_2_ and paO_2_) were monitored at the end of the acquisition to exclude non-physiological conditions. No inter-group differences were observed in mean paCO_2_ (*Shank3B*^*+/+*^: 16.5 ± 4.0 mmHg; *Shank3B*^*-/-*^: 18.6 ± 4.9 mmHg; *p=*0.29) or paO_2_ (*Shank3B*^*+/+*^: 218.3 ± 40.5 mmHg; *Shank3B*^-/-^: 219.5 ± 29.7 mmHg; *p* = 0.94) between mutants and control mice. Possible genotype-dependent differences in anesthesia sensitivity were evaluated with mean arterial blood pressure, amplitude of cortical BOLD signal fluctuations and minimal alveolar concentration, three independent readouts previously shown to be correlated with anesthesia depth (Steffey et al., 2003; Liu et al., 2010; Zhan et al., 2014).

Functional images were acquired with a 7T MRI scanner (Bruker Biospin, Milan) as previously described (Liska et al., 2015), using a 72-mm birdcage transmit coil and a 4-channel solenoid coil for signal reception. For each session, high-resolution anatomical images were acquired with a fast spin echo sequence (repetition time [TR] = 5500 ms, echo time [TE] = 60 ms, matrix 192 × 192, field of view 2 × 2 cm, 24 coronal slices, slice thickness 500 μm). Co-centered single-shot BOLD rsfMRI time series were acquired using an echo planar imaging (EPI) sequence with the following parameters: TR/TE 1200/15 ms, flip angle 30°, matrix 100 × 100, field of view 2 × 2 cm, 24 coronal slices, slice thickness 500 μm for 500 volumes.

### Functional connectivity analyses

Functional connectivity based on rsfMRI is a method to map temporal dependency of spontaneous fluctuations of the BOLD signal between brain regions during no-task condition and is widely used in human clinical (Van Den Heuvel and Pol, 2010) and preclinical (Liska and Gozzi, 2016) studies to describe the macroscale functional organization of brain networks in autism and other neuropsychiatric disorders. Here we employed rsfMRI connectivity to detect putative derangements of functional networks associated to *Shank3B* homozygous mutation.

Before calculating rsfMRI metrics, raw data was preprocessed as described in previous work (Sforazzini et al., 2016; Liska et al., 2017; Michetti et al., 2017). The initial 50 volumes of the time series were removed to allow for T1 equilibration effects. Data were then despiked, motion corrected and spatially registered to a common reference template. Motion traces of head realignment parameters (3 translations + 3 rotations) and mean ventricular signal (corresponding to the averaged BOLD signal within a reference ventricular mask) were used as nuisance covariates and regressed out from each time course. All rsfMRI time series also underwent band-pass filtering to a frequency window of 0.01–0.1 Hz and spatial smoothing with a full width at half maximum of 0.6 mm.

To obtain a data driven identification of the brain regions exhibiting genotype-dependent alterations in functional connectivity, we calculated voxel-wise long-range and local connectivity maps for all mice. Long-range connectivity is a graph-based metric also known as unthresholded weighted degree centrality and defines connectivity as the mean temporal correlation between a given voxel and all other voxels within the brain (Cole et al., 2010). Local connectivity was instead mapped by limiting the measurement of functional coupling within a 6-voxel radius sphere. Pearsons’ correlation scores were first transformed to *z*-scores using Fisher’s *r*-to-*z* transform and then averaged to yield the final connectivity scores.

To corroborate results of voxel-wise connectivity, we calculated connectivity matrices of 170 unilateral anatomical regions (85 right and 85 left) for each mouse and then we performed a *t*-test between genotypes for each edge separately (*t* > 3.5, *p* < 0.001) to identify temporal correlations between regions disrupted in *Shank3B*^-/-^ mice. FWER correction at the network level was performed using 5000 random permutations (*p* < 0.05) as implemented in the network based statistics (Zalesky et al., 2010).

Target regions of long-range connectivity alterations in *Shank3B*^-/-^ mice were mapped using seed-based analysis in regions or volumes of interest (VOIs). As for long range and local connectivity analysis, r-scores were transformed to z-scores using Fisher’s r-to-z transform before statistics. The location of small ROIs of 3×3×1 voxels (shown as small red squares in Fig. 2B-D) was selected based on between-group regional rsfMRI desynchronization observed with unbiased long-range connectivity. Specifically, seeds were placed in the anterior cingulate to map derangements in antero-posterior connectivity within the default mode network, and bilaterally in the striatum to probe impaired cortico-striatal connectivity. Voxel-wise intergroup differences in local and long range connectivity as well as for seed based mapping were assessed using a 2-tailed Student’s *t*-test (|*t*| > 2, p < 0.05) and family-wise error (FWE) cluster-corrected using a cluster threshold of *p* = 0.01 as implemented in FSL. Antero-posterior DMN and cortico-striatal hypo-connectivity were probed by computing VOI-to-seeds correlations. Prefrontal cortex and striatum were employed as VOIs for separate analysis. The location of seeds employed for mapping is indicated in Fig 2B-D. The statistical significance of intergroup effects was quantified using 2-way repeated-measures ANOVA, where seed location (repeated-measure factor) and genotype (between-group factor) were used as variables. Interhemispheric functional connectivity was delineated by computing correlation coefficients of ROI pairs covering major representative anatomical regions. Based on evidence that interhemispheric functional connectivity may be more heterogeneous in ASD patients with respect to age matched typically developing populations (Hahamy et al., 2015), we also tested the presence of “idiosyncratic” interhemispheric connectivity in *Shank3B*^-/-^ mutants with respect to control mice, by measuring cross-subject variability in inter-hemispheric measurements of homotopic synchronization as described by Hahamy and colleagues.

### Behavioral tests

Behavioral testing was carried out two weeks after the imaging sessions. Mice underwent a male-female social interaction test during the light phase, as previously described (Scattoni et al., 2011; Scattoni et al., 2013). An unfamiliar stimulus, i.e. a control female mouse in estrous, was placed into the home-cage of an isolated test male mouse, and social behavior was recorded during a 3-min test session. To measure ultrasound vocalization (USV) recordings, an ultrasonic microphone (Avisoft UltraSoundGate condenser microphone capsule CM16, Avisoft Bioacoustics) was mounted 20 cm above the cage and the USVs were recorded using Avisoft RECORDER software version 3.2. Settings included sampling rate at 250 kHz; format 16 bit. The ultrasonic microphone was sensitive to frequencies between 10 and 180 kHz. For acoustical analysis, recordings were transferred to Avisoft SASLabPro (version 4.40) and a fast Fourier transformation was conducted as previously described (Scattoni et al., 2008). Start times for the video and audio files were synchronized.

Scoring of social investigation and other non-social behaviors was conducted using Noldus Observer 10XT software (Noldus Information Technology) and multiple behavioral responses exhibited by the test mouse were measured: anogenital sniffing (direct contact with the anogenital area), total sniffing (sniffing or snout contact with the flank and head areas), following behaviors, self-grooming (self-cleaning, licking any part of its own body), mounting, rearing up against the wall of the home-cage, rearing, digging in the bedding and immobility. Social investigation was defined as the sum of total sniffing and following behaviors (Scattoni et al., 2008).

### Structural MRI

To identify putative gray matter reorganization coincident with functional hypo-connectivity in the cortex of *Shank3B*^-/-^ mice, we carried out *post-mortem* Voxel-Based Morphometry (VBM) (Pagani et al., 2016b). Images were acquired on *ex-vivo* in paraformaldehyde fixed specimens, a procedure employed to obtain high-resolution images with negligible confounding contributions from physiological or motion artefacts. Brains were imaged inside intact skulls to avoid post-extraction deformations. *Shank3B*^-/-^ and control mice were deeply anaesthetized with an Isoflurane 5% and their brains were perfused *in situ* via cardiac perfusion (Dodero et al., 2013; Sannino et al., 2013). The perfusion was performed with phosphate buffered saline followed by paraformaldehyde (4% PFA; 100 ml, Sigma, Milan). Both perfusion solutions were added with a Gadolinium chelate (Prohance, Bracco, Milan) at a concentration of 10 and 5 mM, respectively, to shorten longitudinal relaxation times. High-resolution morpho-anatomical T2-weighted MR imaging of mouse brains was performed using a 72 mm birdcage transmit coil, a custom-built saddle-shaped solenoid coil for signal reception. For each session, high-resolution morpho-anatomical images were acquired with the following imaging parameters: FLASH 3D sequence with TR = 17 ms, TE = 10 ms, α = 30°, matrix size of 260 × 180 × 180, field of view of 1.83 × 1.26 × 1.26 cm and voxel size of 70 μm (isotropic).

### Voxel-Based Morphometry (VBM) of Gray Matter (GM) Volume

Inter-group morpho-anatomical differences in local GM volumes were mapped using a registration-based VBM procedure (Pagani et al., 2016a; Pagani et al., 2016b). Briefly, high-resolution T2-weighted images were corrected for intensity non-uniformity, skull stripped and spatially normalized to a study-based template using affine and diffeomorphic registrations. Registered images were segmented to calculate tissue probability maps. The separation of the different tissues is improved by initializing the process with the probability maps of the study-based template previously segmented. The Jacobian determinants of the deformation field were extracted and applied to modulate the GM probability maps calculated during the segmentation. This procedure allowed the analysis of GM probability maps in terms of local volumetric variation instead of tissue density. Brains were also normalized by the total intracranial volume to further eliminate overall brain volume variations and smoothed using a Gaussian kernel with a sigma of three-voxel width.

For statistical analysis, voxel-wise intergroup differences in regional GM volume differences between *Shank3B*^-/-^ mice and controls were mapped using a 2-tailed Student’s *t*-test (|*t*| > 2, *p* < 0.05) and FWE cluster-corrected using a cluster threshold of *p* = 0.01 (Worsley et al., 1992) as implemented in FSL (Jenkinson et al., 2012). To quantify volumetric changes identified with VBM, we employed preprocessed images to independently calculate the size of neuroanatomical areas via volumetric anatomical labelling (Pagani et al., 2016b).

### Diffusion MRI

Fixed brains also underwent DW imaging. Each DW data set was composed of 8 non-DW images and 81 different diffusion gradient-encoding directions with *b* = 3000 s/mm^2^ (∂ = 6 ms, Δ = 13 ms) acquired using an EPI sequence with the following parameters: TR/TE = 13500/27.6 ms, field of view 1.68 × 1.54 cm, matrix 120 × 110, in-plane spatial resolution 140 × 140 μm, 54 coronal slices, slice thickness 280 μm, number of averages 20 as recently described (Dodero et al., 2013; Liska et al., 2017).

### White matter fiber tractography and tract-based spatial statistics

The DW datasets were first corrected for eddy current distortions and skull-stripped. The resulting individual brain masks were manually corrected using ITK-SNAP (Yushkevich et al., 2006). Whole brain tractography was performed with MRtrix3 (Tournier et al., 2012) using constrained spherical deconvolution (lmax = 8) and probabilistic tracking (iFOD2) with a FOD amplitude cut-off of 0.2. For each dataset, the whole brain mask was used as a seed, and a total of 100,000 streamlines were generated. The corpus callosum and cingulum were selected as tracts of interest, given their major cortico-cortical extension and direct involvement in prefrontal-posterior connectivity. The tracts were virtually dissected with waypoint VOIs previously described (Liska et al., 2017) using TrackVis (http://www.trackvis.org/). Inter-group differences in streamline counts of the tracts were evaluated using a two-tailed Student’s *t*-test (*t* > 2, *p* < 0.05). For visualization purposes, dissected corpus callosum and cingulum were transformed to the Allen Mouse Common Coordinate Framework, version 3 (http://www.brain-map.org/). Further, we performed a Tract-Based Spatial Statistics (TBSS) analysis, as implemented in FSL (Smith et al., 2006) and previously described (Dodero et al., 2013). Fractional anisotropy (FA) maps from all subjects were non-linearly registered to an in-house FA template with FSL-FLIRT and FSL-FNIRT and thinned using a FA threshold of 0.2 to create a skeleton of the white matter. Voxel-wise inter-group differences between deletion and control mice were evaluated with permutation testing using 5000 permutations (*p* < 0.05) (Winkler et al., 2014). FA was also quantified and compared between *Shank3B*^-/-^ mice and controls for genotype dependent alterations in representative regions, such as corpus callosum, dorsal hippocampal commissure and forceps minor using a 2-tailed Student’s *t*-test (*p* < 0.05)

### Virus production and injection

Unpseudotyped recombinant SADΔG-mCherry Rabies Virus was produced as described by Bertero and colleagues, 2018. Briefly, B7GG packaging cells, which express the rabies envelope G protein, were infected with unpseudotyped SADΔG-mCherry-RV, obtained by courtesy of Prof. Edward Callaway. Five to six days after infection, viral particles were collected, filtrated through 0.45 μm filter and concentrated by two rounds of ultracentrifugation. The titer of the SADΔG-mCherry-RV preparation was established by infecting Hek-293T cells (ATCC cat n° CRL-11268) with tenfold serial dilution of viral stock, counting mCherry expressing cells 3 days after infection. The titer was calculated as 2×10^11^ Infective Units/ml (IU/ml), and the stock was therefore considered suitable for in vivo microinjection. Mice were anesthetized with isoflurane (4%) and firmly stabilized on a stereotaxic apparatus (Stoelting Inc.). A micro drill (Cellpoint Scientific Inc.) was used to drill holes through the skull. RV injections were carried out as previously described in adult (12–16-weeks-old) male *Shank3B*^*-/-*^ and control *Shank3B*^*+/+*^ littermates (Cavaccini et al., 2018). Injections were performed with a Nanofil syringe mounted on an UltraMicroPump UMP3 with a four channel Micro4 controller (World Precision Instruments), at a speed of 5 nl per seconds, followed by a 5-10 minutes waiting period, to avoid backflow of viral solution and unspecific labelling. One microliter of viral stock solution was injected unilaterally in the primary cingulate cortex of 10-20 weeks old *Shank3B*^-/-^ and wild-type control mice. Coordinates for injections, in mm from Bregma: +1.42 from anterior to posterior, +0.3 lateral, −1.6 deep.

### Immunohistochemistry and image analysis

Seven days after RV injection, animals were transcardially perfused with 4% paraformaldehyde (PFA) under deep isoflurane anesthesia (5%), brains were dissected, post-fixed over night at 4°C and vibratome-cut (Leica Microsystems). RV infected cells were detected by means of immunohistochemistry performed on every other 100 μm thick coronal section, using rabbit anti Red-Fluorescent-Protein (RFP) primary antibody (1:500 AbCam), and Goat anti Rabbit-HRP secondary antibody (1:500 Jackson immunoresearch), followed by 3-3’ diaminobenzidine tetrahydrochloride (DAB, Sigma Aldrich) staining. Wide-field imaging was performed with a MacroFluo microscope (Leica) and RGB pictures where acquired at 1024×1024 pixel resolution. Labelled neuron identification was manually carried out in n = 4 *Shank3B*^*-/-*^ and n=5 *Shank3B*^*+/+*^ control littermates by an operator blind to the genotype, while the analysis was performed using custom made scripts to automatically register each brain section on the corresponding table on the Allen Brain Atlas (http://www.brain-map.org) and to count labelled neurons and assign them to their anatomical localization. Final region-based cell population counts were expressed as fraction of the total amount of labeled cells counted per animal.

### NeuN+ cell density quantification

Six anatomically comparable 100 μm thick coronal sections of *Shank3B*^-/-^ and control littermates (N = 5, each) were processed for NeuN immunostaining and neural density quantification in the anterior cingulate cortex. Immunofluorescence on free floating sections was performed using mouse anti NeuN primary antibody (1:1000 Millipore, #MAB377), overnight, followed by 2 hours rhodamine-red goat anti mouse secondary antibody (1:500 Invitrogen, R6393). Images were acquired on a Nikon A1 confocal system, equipped with 561 diode. Z series of 5 stacks (1μm z-step) confocal images, using a 10x plan-apochromat objective, were acquired at 1024×1024 pixel resolution. NeuN positive cells in the anterior ngulate cortex were manually counted by an operator blind to the genotype, using the Cell Counter plugin of Fiji (Schindelin et al., 2012). Cell count for each section was normalized on the number of pixel in the area. Data are expressed as mean number of NeuN+ cells every 100 pixels per genotype.

## Supplementary figures

**Figure S1.**
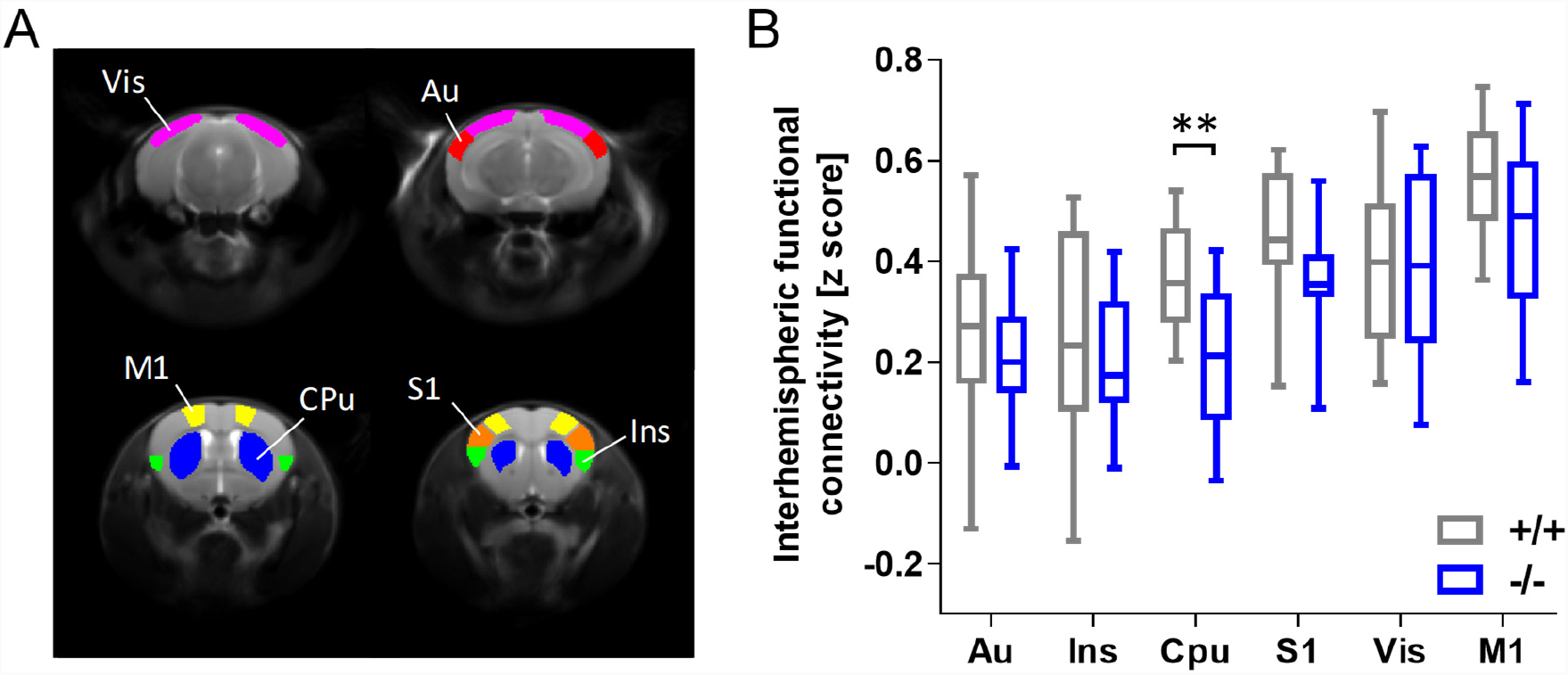
Reduced striatal inter-hemispheric connectivity in *Shank3B*^-/-^ mice. Inter-hemispheric rsfMRI connectivity calculated between time courses extracted from representative VOIs depicted in (A). The resulting inter-hemispheric *r*-scores were transformed to *z*-scores using Fisher’s *r*-to-*z* transform (B). Au: auditory cortex; Ins: insular cortex; CPu: striatum; S1: primary somatosensory cortex; Vis: visual cortex; M1: motor cortex. \*\**p* < 0.01, Student’s *t*-test.

**Figure S2.**
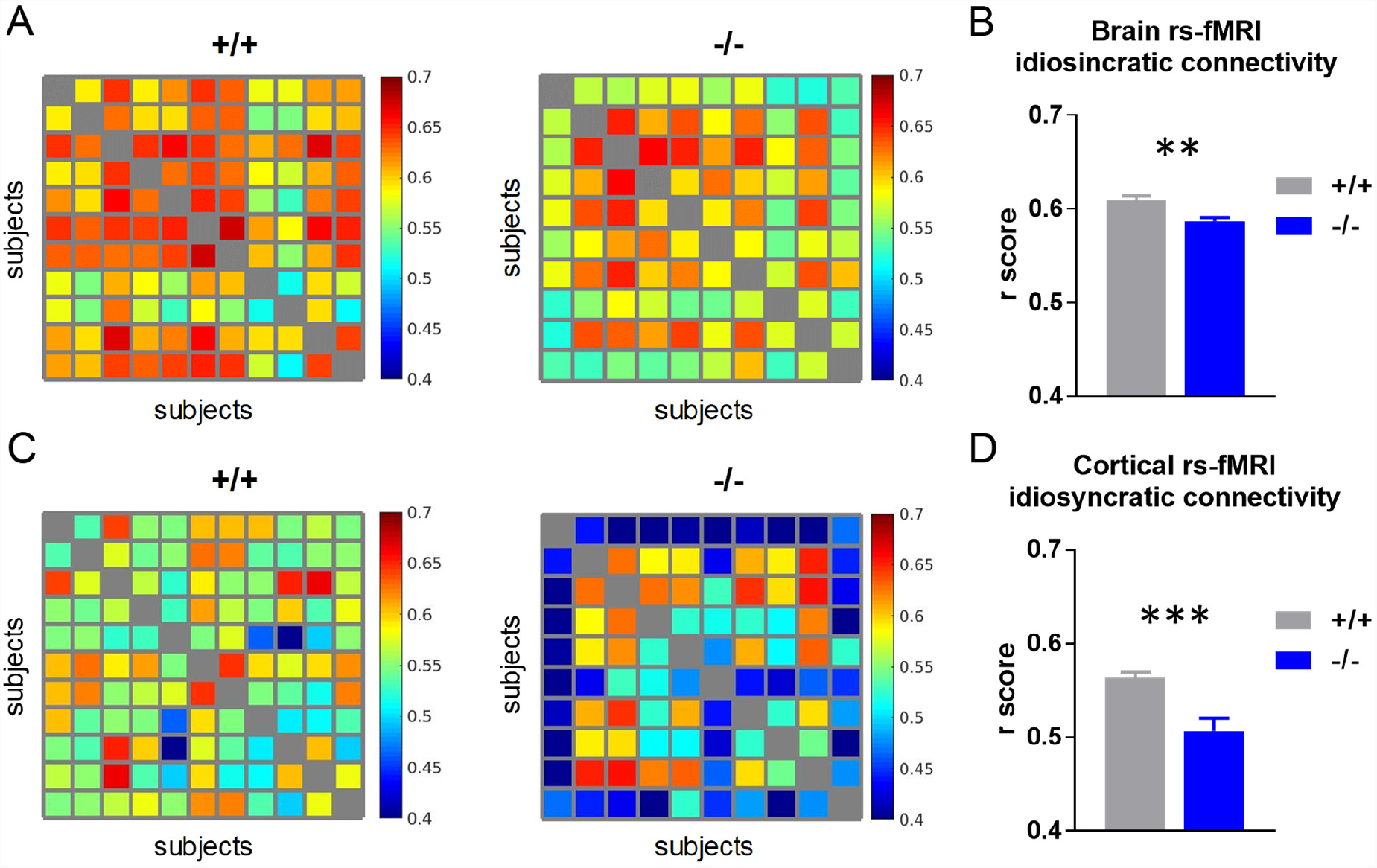
Idiosyncratic inter-hemispheric connectivity in *Shank3B*^-/-^ mice. Each cell represents the inter-subject Pearson’s correlation value between a pair of homotopic inter-hemispheric connectivity maps. Homotopic connectivity maps were calculated voxel-wise for each mouse (A) considering the whole brain and (C) the cortex only. (B) Decreased inter-subject similarity in the Shank3B^-/-^ cohort compared to control mice (*t*-test, *t*_98_ = 2.98, *p* = 0.004). (D) Idiosyncratic homotopic connectivity was particularly apparent in the neocortex *Shank3B*^-/-^ mice (*t*-test, *t*_98_ = 3.93, *p* < 0.001). \*\**p* < 0.01, \*\*\**p* < 0.001.

**Figure S3.**
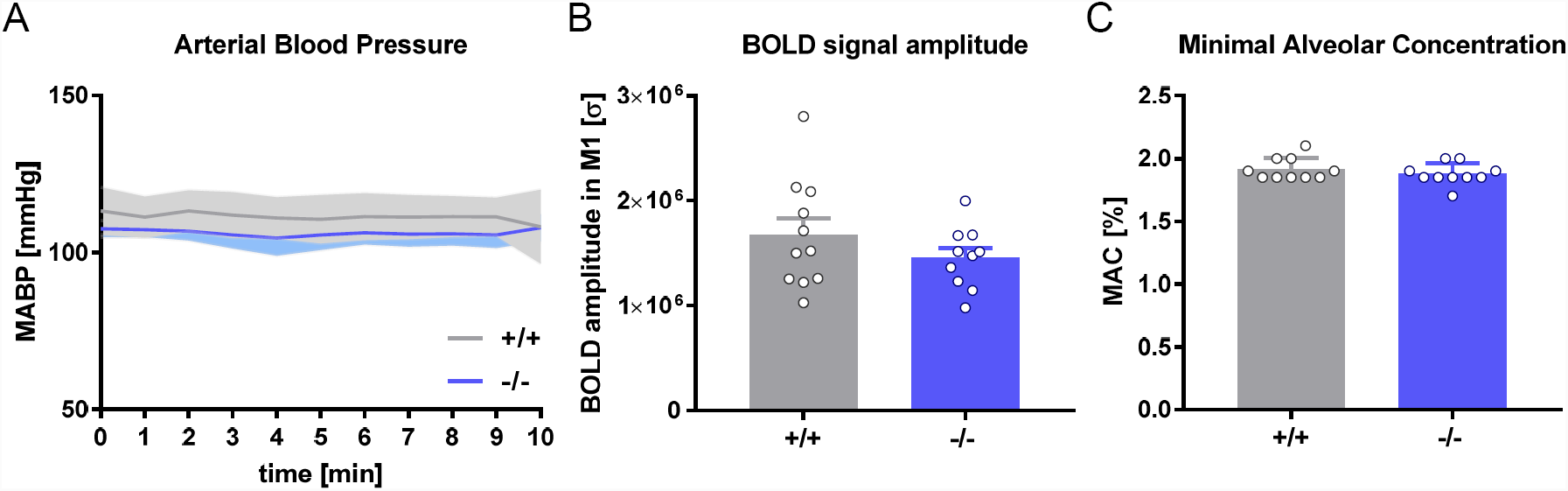
Measurements of anesthesia sensitivity. Statistical testing did not reveal any genotype-dependent differences in anesthesia sensitivity as measured with (A) mean arterial blood pressure mapping, (B) amplitude of cortical BOLD signal fluctuations as measured in a representative region (M1) and (C) minimal alveolar concentration (Student’s *t*-test). M1: primary motor cortex.

**Figure S4.**
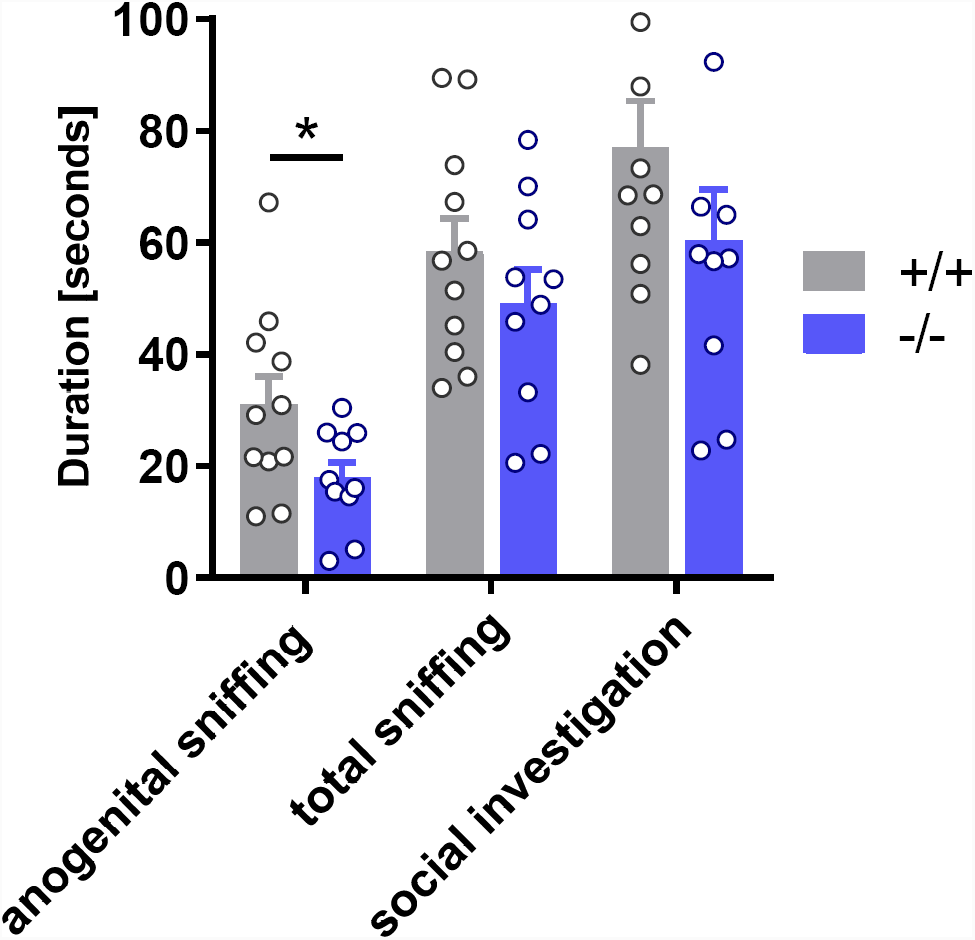
Social investigation in *Shank3B*^-/-^ mice. Anogenital sniffing (*t*-test, *t*_19_ = 2.211, *p=* 0.04), total sniffing (*t*-test, *t*_19_ = 1.09, *p*= 0.29) and social investigation (*t*-test, *t*_19_ = 1.34, *p*= 0.19) in *Shank3B*^-/-^ mutants and *Shank3B*^+/+^ control mice. \**p* < 0.05, Student’s *t*-test, uncorrected.

